# Nutrient requirements for cell differentiation progression in *Batrachochytrium dendrobatidis*

**DOI:** 10.1101/2025.05.28.656562

**Authors:** Naomi C. Okada, Tre’Shur Williams-Carter, Joelyne Contreras, Trae Hill, Dajahneek O’Brien, Ernesto Abel-Santos

## Abstract

*Batrachochytrium dendrobatidis* (*Bd*) is the causative agent of the deadly amphibian disease chytridiomycosis, which has decimated amphibian populations across the globe. *Bd* reproduces via a multistage asexual pathway. The *Bd* life cycle begins with a motile zoospore. Upon contact with a surface, the zoospore anchors itself and forms a cyst. The cyst then develops into a germling that produces rhizoid structures. Finally, the germling matures into a zoosporangium which then releases multiple new zoospores.

In this study, we investigated the effects of nutrients on the growth and differentiation of *Bd*. The presence of carbohydrates, vitamins, and trace minerals had a negligible effect on *Bd’s* ability to progress through its life cycle. In contrast, the amino acids L-leucine and L-isoleucine (and to a lesser extent L-arginine) were required for cysts to germinate into germlings. However, none of these amino acids were able to sustain further cellular development. Furthermore, the presence of nitrate or ammonium aided progression from germlings into immature zoosporangia that were not able to produce zoospores. The progression to mature zoosporangia was blocked by nitrite or the combined action of calcium ions and bicarbonate. The addition of soluble and insoluble protein sources alone also allowed *Bd* to develop into immature zoosporangia and affected the rate of development. Finally, protein sources in combination with L-amino acids allowed the completion of the *Bd* cycle by allowing development from zoospores to mature zoosporangia that were able to release new zoospores.

**Importance:** Even though the cell differentiation cycle of *Bd* has been microscopically described, not much is known about the requirements for *Bd* to progress throughout its life stages. The identification of the molecular cues that allow *Bd* cell cycle progression may be crucial to disease mitigation efforts. As a corollary, we have identified conditions that allow for the arrest of *Bd* at various stages of its life cycle. By changing the metabolite availability, we can efficiently obtain cultures enriched for zoospores, cysts, germlings, immature zoosporangia, or spore-releasing zoosporangia. The ability to temporarily freeze and then restart *Bd* at different points of its life cycle, can allow for determining the mechanisms behind individual transitions between cell types.

## Background

In the late 1990’s, multiple studies described an unknown chytrid fungus on skin samples of deceased amphibians in the Americas and Australia^1–3^. This novel species was dubbed *Batrachochytrium dendrobatidis*^2^ (*Bd*) and was later established to cause the fatal skin disease chytridiomycosis^4^.

Since its discovery, *Bd* has been identified in over 1,000 species of amphibians across all continents^5^. Furthermore, *Bd* and its sister species *Batrachochytrium salamandrivorans* (*Bsal*) have been implicated in the extinction or decline of at least 6.5% of all known amphibian species worldwide. Indeed, amphibian chytridiomycosis has been described as the pathogenic disease associated with the greatest known loss of vertebrate biodiversity^6^.

While protocols have been developed to treat and clear chytridiomycosis in captive amphibians, antifungal agents tend to have narrow therapeutic windows or toxic effects on the hosts^7^. Furthermore, antifungal agents do not seem to decrease zoospore load in either laboratory or field tests^8,9^.

*Bd* undergoes cell differentiation that allows it to survive in the environment and invade the amphibian host. The complete asexual reproductive cycle of *Bd* consists of two main stages: a small (generally 3-5 μm in diameter) motile zoospore and a larger sessile thallus^2^. The flagellated zoospores will attach and encysting onto a solid surface^10^. In the absence of any nutrients, cysts will remain viable but will not complete their lifecycle^11^. In contrast, zoospores that encyst with adequate nutrients develop into germlings with thread-like rhizoids^2^. These germlings will then grow into mature thalli containing one or more compartments called zoosporangia. Multiple new zoospores form within each of these zoosporangia and are released via a discharge papilla to complete *Bd’s* life cycle.

Prevalence of *Bd* within a population of amphibians is negatively correlated with population size for at least some species^12^. Additionally, the number of zoospores that an individual amphibian is exposed to is negatively correlated to amphibian survival^13^. Therefore, a thorough understanding of the ability of *Bd* to differentiate may be crucial for disease mitigation.

While the epidemiology of *Bd* is well-studied, relatively little is known about *Bd*’s cellular biology, including molecular mechanisms behind host cell invasion^14^ or fungal development^15,16^. In this study, we determined the nutritional signals required for *Bd* to complete each stage of its life cycle. We found that only hydrophobic amino acids, inorganic nitrogen, and protein sources were necessary for *Bd* differentiation. We also found that by limiting nutrient availability, *Bd* differentiation can be arrested at the cyst, germling, or zoosporangia stages, while maintaining cell viability.

## Materials and Methods

### Reagents and materials

Amino acids, lactose monohydrate, D-(+)-mannose, D-(+)-galactose, riboflavin, folic acid, nicotinic acid, pyridoxine, 4-aminobenzoic acid (PABA), calcium pantothenate, biotin, MoNa_2_O_4_, KCHO_3_, NH_4_Cl, and Na_2_SeO_3_ were obtained from Sigma-Aldrich (St. Louis, MO, USA); agar and tryptone (pancreatic digest of casein) were obtained from BD (Franklin Lakes, NJ, USA); gelatin hydrolysate was obtained from EMD Millipore (Burlington, MA, USA); cell culture-grade water and D-glucose were obtained from VWR (Radnor, PA, USA); 10,000 I.U./mL Cellgro Penicillin Streptomycin Solution was obtained from Corning (Corning, NY, USA); dimethyl sulfoxide (DMSO) was obtained from Biotium (San Francisco, CA, USA); fetal bovine serum was obtained from Gibco (Waltham, MA, USA); MgCl_2_ was obtained from Research Organics (Cleveland, OH, USA); KH_2_PO_4_, CaCl_2_, ZnCl_2_, and CoCl_2_ were obtained from Alfa Aesar (Haverhill, MA,USA); NH_4_CHO_3_ and L-arabinose were obtained from Fluka (Seelze, Germany); Ca(NO_3_)_2_ was obtained from Mallinckrodt Chemicals (Staines-upon-Thames, UK); KNO_3_ and KNO_2_ were obtained from Ward’s Science (Rochester, NY, USA); D-xylose was obtained from Fisher Chemical (Pittsburgh, PA, USA); D-trehalose and sucrose were obtained from MP Biomedicals (Santa Ana, CA, USA); D-raffinose pentahydrate was obtained from Amresco (Solon, OH, USA); L-rhamnose was obtained from Difco (Franklin Lakes, NJ, USA); D-(-)-fructose and MnSO_4_ were obtained from J.T. Baker (Phillipsburg, NJ, USA); CuSO_4_ and bovine serum albumin (BSA) were obtained from Acros Organics (Geel, Belgium); NH_4_VO_3_ was obtained from Strem Chemicals (Newburyport, MA, USA); porcine gelatin was obtained from GBiosciences (St. Louis, MO, USA); carnitine was obtained from TCI (Montgomeryville, PA, USA); and FeCl_3_·6H_2_O was obtained from BDH (San Jose, CA, USA) Representative lizard scales and bird feathers were collected as natural shedding from captive *Pogona vitticeps* and *Columba livia*, respectively. Fish (*Oncorhynchus nerka*) scales were obtained from Costco (Henderson, NV, USA). Frog skin from *Lithobates onca*, and *Agalychnis callidryas* were harvested from available carcasses. All keratin sources were autoclaved in deionized water (DI-H_2_O) prior to introduction to cell cultures.

### Acquisition and culturing of Bd

*Bd* strain LBP114 was isolated from an infected *Lithobates onca* individual (collected on March 16, 2017 from Blue Point Springs, Lake Mead, NV) by the Jef Jaegar group (UNLV). *Bd* cultures were maintained in TGhL broth (1.6% (w/v) tryptone, 0.4% (w/v) gelatin hydrolysate, 0.2% (w/v) lactose)^17^ or H-broth (1% (w/v) tryptone, 0.32% (w/v) D-glucose)^18^ at 4 °C and sub-cultured into fresh media every 8 weeks. For each passage, 1 ml of liquid culture was spread onto TGhL agar (1.6% (w/v) tryptone, 0.4% (w/v) gelatin hydrolysate, 0.2% (w/v) lactose, 1% (w/v) agar, 0.2 U/ml penicillin-streptomycin) plates^18^ and incubated for 4-5 days at 19 °C. After incubation, a small section of agar was transferred into 25-30 ml of fresh H-broth and grown for at least 1 week at 19 °C and then stored at 4 °C before further use.

Samples were also cryo-archived at −80 °C, as described previously^19^. Briefly, H-broth aliquots containing zoospores and zoosporangia were pelleted and resuspended in cryopreservation media (TGhL broth supplemented with 10% DMSO and 10% fetal bovine serum). The mixed cell population was aliquoted into 2-ml cryotubes, and frozen in a cryo-freezing container at −80 °C. To revive cell cultures, cryotubes were briefly thawed in a 43 °C water bath, plated at high density onto TGhL plates, and incubated for 4-5 days at 19 °C. After incubation, a small section of agar was transferred into 25-30 ml of fresh H-broth and grown for at least 1 week at 19 °C and then stored at 4°C before further use.

### Zoospore purification

A 1-2 ml aliquot of a *Bd* mixed liquid culture was spread onto TGhL agar plates and incubated for 4-5 days at 19 °C. After incubation, the plates were flooded with 4 °C sterile water and incubated at 4 °C for at least 30 minutes. Plates were then harvested with a plastic cell scraper and washed with additional sterile water. The collected samples were pooled, and vacuum filtered through a Whatman 2 or a Whatman 3 filter paper. The filtrate was then centrifuged at 1,750 ×g for 5 minutes. Pellets were resuspended in a 2X concentration of dilute salt solution (DSS) (2 mM KH_2_PO_4_, 0.2 mM MgCl_2_ and 40 μM CaCl_2_)^20^, re-pelleted, and finally resuspended in fresh 2X DSS. Cell density was adjusted with additional fresh 2X DSS to achieve an optical density at 492 nm (OD_492_) of between 0.012-0.024. Microscopy observation showed that the resulting zoospore suspensions were more than 95% pure.

### Testing nutrient media

Aliquots (25 μl) of purified zoospore suspensions were added into the 60 central wells (B2 through G11) of a 96-well plate. The outside wells were filled with autoclaved water to prevent desiccation of the internal wells. Zoospore suspensions were supplemented with either 25 μl of experimental nutrient media, positive (2X H-broth) controls, or negative (sterile DI-H_2_O) controls.

Nutrient solutions used for these experiments started with a minimal defined medium (MDM) based on previously published literature^21,22^. The final composition of the MDM was: (*i*) The twenty proteinaceous amino acids (0.2 mg/ml each); (*ii*) Ca(NO_3_)_2_ (10 mM) and NH_4_HCO_3_ (20 mM) as inorganic nitrogen sources; (*iii*) D-glucose (80 μg/ml) as a carbon source; (*iv*) pyridoxine, folic acid, riboflavin, pantothenic acid, thiamine, nicotinic acid, biotin, para-aminobenzoic acid (PABA), and carnitine as vitamins (1 μg/ml each); and (*v*) CuSO_4_ (1 μg/ml), MoNa_2_O_4_ (0.4 μg/ml), MnSO_4_ (3 μg/ml), ZnCl_2_ (4 μg/ml), CoCl_2_ (50 ng/ml), FeCl_3_ (2 ng/ml), Na_2_SeO_4_ (0.2 μg/ml), and NH_4_VO_3_ (6 ng/ml) as trace minerals. All nutrient solutions were pH balanced to 6.0-7.0 and filter sterilized.

To test for nutritional requirements, MDM was individually modified as needed to include or omit different components. For the amino acid experiments, MDM was modified to omit one group of structurally related amino acids: small (L-Ala, Gly, L-Ser, L-Thr, L-Cys), hydrophobic (L-Leu, L-Ile, L-Val, L-Met), aromatic (L-Phe, L-Tyr, L-Trp), basic (L-Arg, L-Lys, L-His), acidic (L-Asp, L-Glu), amide (L-Asn, L-Gln), or constrained (L-Pro).

For the inorganic nitrogen experiment, Ca(NO_3_)_2_ and NH_4_CHO_3_ in MDM were substituted by Ca(NO_3_)_2_ alone, NH_4_CHO_3_ alone, NH_4_CHO_3_ and KNO_3_, NH_4_CHO_3_ and KNO_2_, NH_4_CHO_3_ and CaCl_2_, KCHO_3_ and Ca(NO_3_)_2_, or NH_4_Cl and Ca(NO_3_)_2_.

For the carbohydrate experiment, glucose in MDM was replaced by D-mannose, D-xylose, D-fructose, D-raffinose, L-rhamnose, L-arabinose, sucrose, α-lactose, D-galactose, or D-trehalose. For the vitamin experiment, vitamins (pyridoxine, folic acid, riboflavin, pantothenic acid, thiamine, nicotinic acid, biotin, PABA or carnitine) were individually omitted from MDM. For the trace mineral experiment, minerals (CuSO_4_, MoNa_2_O_4_, MnSO_4_, ZnCl_2_, CoCl_2_, FeCl_3_, Na_2_SeO_4_, or NH_4_VO_3_) were individually omitted from MDM.

For the protein sources experiment, autoclaved keratin tissues (*P. vitticeps* scales, *C. livia* feathers, *O. nerka* scales, *L. onca* skin, and *A. callidryas* skin) or filter sterilized soluble proteins (BSA or soluble gelatin fraction) were introduced to MDM or DSS, as specified.

### Viability of nutrient-deprived Bd

Freshly harvested zoospores were incubated in DSS for increasing amounts of time. Each day, three separate wells containing *Bd* in DSS were supplemented with H-broth. Resumption of growth was determined by increases of signal at OD_492_.

### Imaging and measurement

The 96-well plates containing *Bd* growth were imaged at specific days following incubation at 19 °C. The images were captured using a Nikon Eclipse TE200-U microscope equipped with a CoolSNAP Dyno camera using a 40X objective lens under brightfield lighting. Each image represented a single random field of view of a given sample within each well, and each well was imaged once. Images were saved as .ND2 files. Each image was visually examined to determine the final stage of cellular development progression under each condition tested.

To measure the area of each cell, images were opened in Fiji^23^ using the Bio-Formats plugin^24^. Regions of interest (ROIs) were then manually drawn around all cells within the field of view using the circular selection tool. The areas of each ROI were measured using Fiji’s built-in Measurement tool. In Microsoft Excel, bins were defined depending on the maximum extent of development visually observed within each treatment group (**Figure 1**). All analyses included bins for zoospores (diameter less than 5.0 μm) and cysts (diameter greater than 5.0 μm). If rhizoid formation was observed, cysts were restricted to particles with diameters between 5.0 μm and 6.0 μm, and a third bin for germlings (greater than 6.0 μm) was included. If the beginnings of internal structures were observed, then germlings were restricted to particles with diameters between 6.0 μm and 8.8 μm, and a fourth bin for zoosporangia (diameter greater than 8.8 μm) was also included. Finally, under some conditions, zoosporangia failed to produce and release new zoospores. Hence, we distinguished between treatment groups that were able to support the development of immature zoosporangia (those that did not form zoospores) from mature zoosporangia (those that formed and released new zoospores), even though both structures showed similar size.

**Figure 1.**
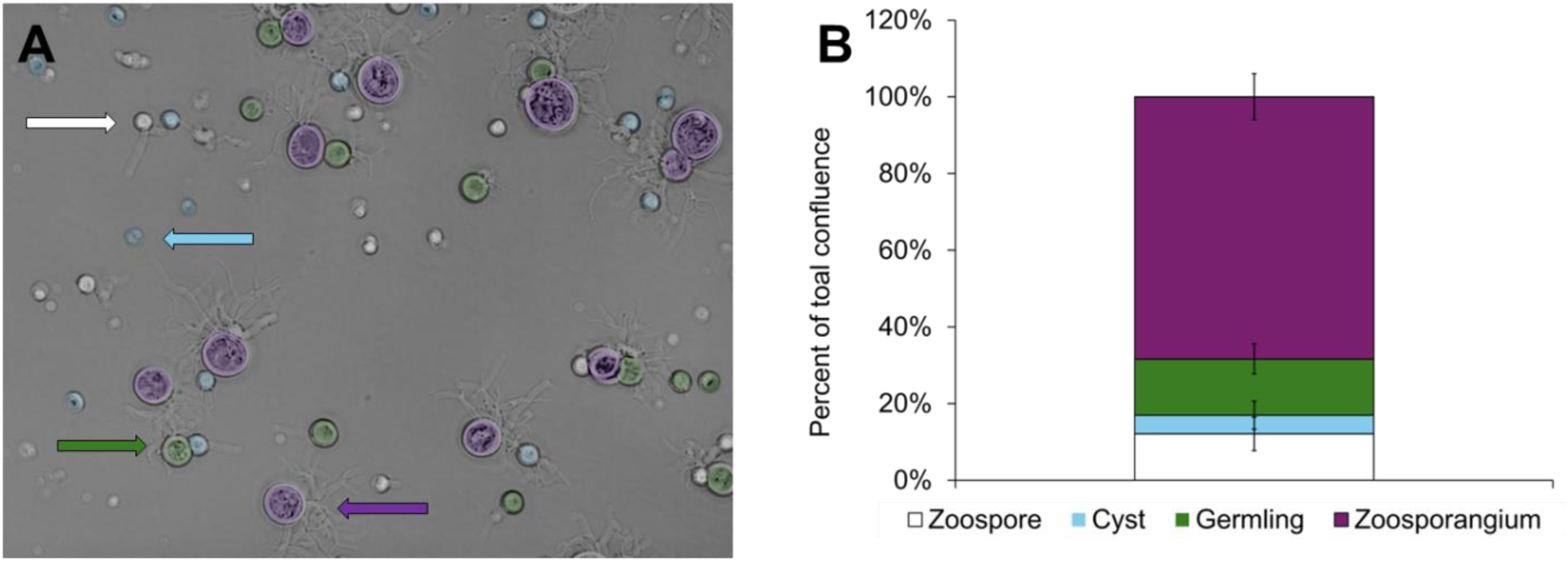
Sample binning: **(A)** A representative image of cells incubated for 3 days in H-broth was analyzed and constituent cells were assigned into bins: a zoospore bin (white; diameter less than 5.0 μm), a cyst bin (light blue; diameter between 5.0 μm and 6.0 μm), a germling bin (green; diameter between 6.0 μm and 8.8 μm) with rhizoids, or a zoosporangium bin (purple; diameter greater than 8.8 μm) with developing internal structures. **(B)** A graphical representation of the data obtained from panel A. Percent of total confluence was calculated by adding the individual area covered by each cell type. The total area for each cell type was then divided by the total area covered by all cell types. For example, to calculate the percent of total confluence of zoosporangia, we added the areas of all purple zoosporangia in panel A. We then divided this value by the sum of the areas of all white zoospores, all light blue cysts, all green germlings, and all purple zoosporangia. Error bars illustrate the standard deviation among all replicates for a given treatment. The error bars shown in this figure are arbitrary representation for the purposes of illustration.

To quantify the proportions of cell types in each experimental population, we calculated the total area covered by an individual cell type. Individual cell type total area was then divided by the total area of each image covered by all cell types. This ratio was designated as percent total confluence and served to quantitatively measure growth and development under the tested nutrient conditions. The appropriateness of percent confluence as a quantitative measure of cell type proportions was validated by comparing the obtained values with visually counting the number of each cell type on selected microscopic fields (**Figure 1**)

### Statistical analysis

Each experiment was performed with six experimental replicates. Results from different conditions were compared using one-way ANOVA followed by standard Tukey *post-hoc* analysis with a 95% confidence interval using GraphPad Prism version 10.2.3 for Windows (GraphPad Software, Boston, Massachusetts USA, www.graphpad.com). Averages and standard deviations were plotted using Microsoft Excel.

## Results

### Quantification of Bd cellular stages

Because the population of *Bd* cell stages strongly favored cells with smaller sizes (*e.g.*, the ratio of individual zoosporangia to zoospores was extremely low), we found that manually counting cell numbers to be inaccurate. This was to be expected because most zoospores do not fully mature into zoosporangia, especially when in low-nutrient conditions and when isolated from other life stages^2^. However, while earlier cell stages eclipsed later cell stages by cell count, the larger size of the later cell stages allowed them to cover a larger surface area. Hence, the percentage of the area (percent total confluence) covered by each cell type provided a better approximation to their relative proportions in our experiments.

Based on previous work^25^ and our microscopic measurements, we created diameter cutoffs for each cell type during *Bd* differentiation. Because of overlap in their diameters, size measurements for germlings and zoosporangia, were supplemented with visual detection of rhizoid structures (for germlings) and internal structures (for zoosporangia). The combination of size measurements and visual structure detection allowed us to enumerate proportions of cell types in each experimental population (**Figure 1**).

### Effect of complex media on Bd differentiation

As previously reported^26^, the vast majority of *Bd* zoospores incubated without nutrients (in DSS) showed no immediate physical changes (**Figure 2A**). A few zoospores (<10%) were able to encyst and arrested at this stage (**Figure 2B**). Prolonged incubation in DSS resulted in shriveled cysts. The arrested cysts were nonetheless viable for at least two seeks and continued their life cycle when nutrients were reintroduced (**Figure S1**).

**Figure 2.**
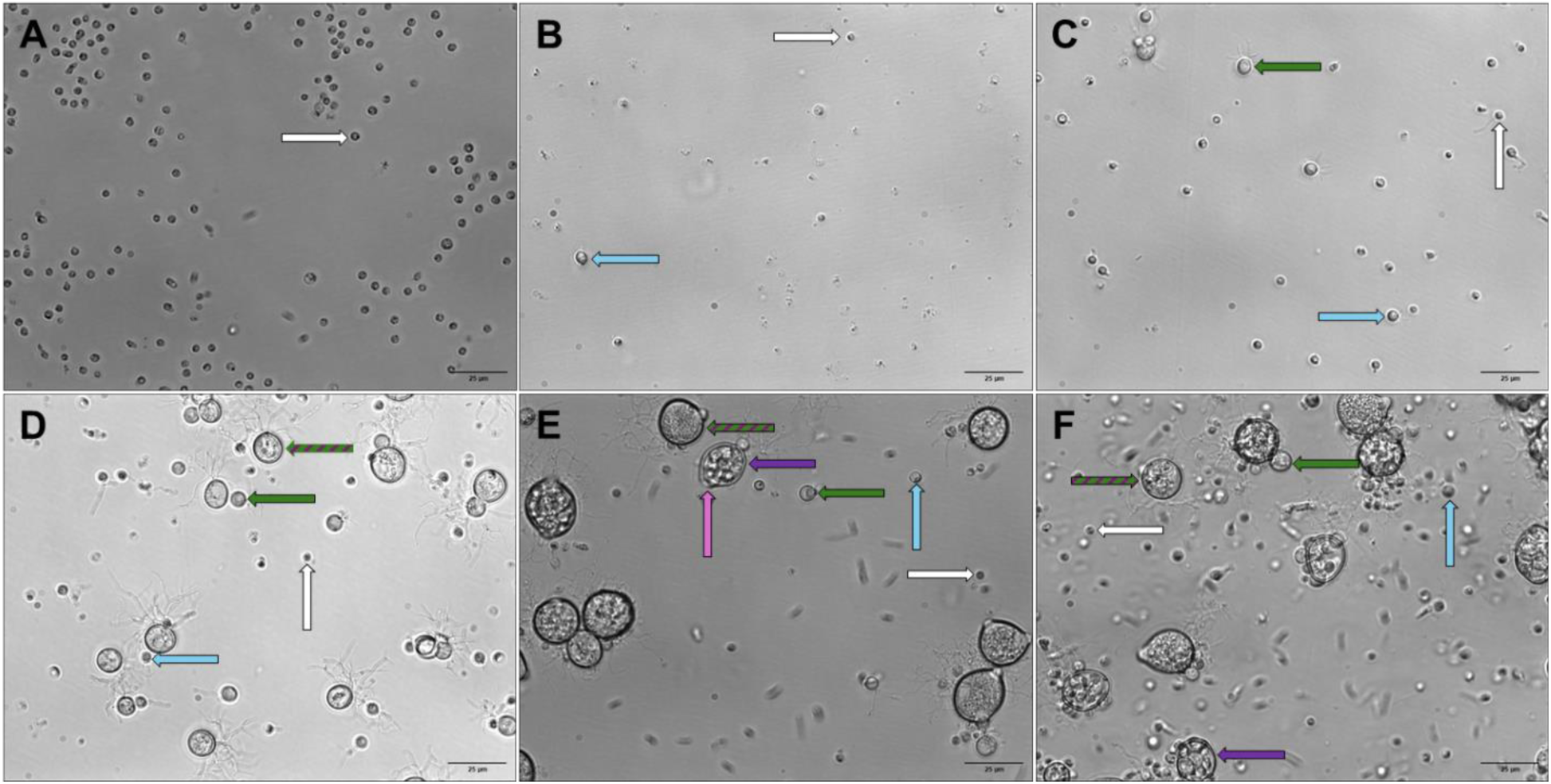
*Bd* cell cycle stages. *Bd* zoospores were incubated at 19 °C for **(A)** 1 hour or **(B)** for 1 day in DSS, or for **(C)** 1 day, **(D)** 2 days, **(E)** 4 days, or **(F)** 5 days in H-broth. Representative zoospores (white arrows), cysts (blue arrows), germlings (green arrows), immature zoosporangia (purple arrows with green stripes), mature zoosporangia (purple arrows), and discharge papilla (magenta arrow) are indicated.

In contrast, zoospores incubated in media containing carbon and nitrogen sources (H-broth) were able to complete the entire *Bd* life cycle. After 1 day incubation in H-broth, approximately 40% of the *Bd* population consisted of cysts (**Figure 2C**). By day 2, most cysts had progressed to the germling stage, and a few immature zoosporangia appeared (**Figure 2D**). By days 3-4, zoosporangia became the dominant cellular form (**Figure 2E**). On days 4-5 of incubation, new zoospores were released from the mature zoosporangia and the new zoospores started to encyst (**Figure 2F**). After this point, the *Bd* population in H-broth became too overgrown to measure accurately. To quantify these changes, we converted the data to a graphic form as depicted in **Figure 1A** to **Figure 1B**.

The minimal defined medium (MDM) contains a combination of proteinaceous amino acids, inorganic nitrogen sources, D-glucose, vitamins, and trace minerals at pre-calculated concentrations. In contrast to DSS, the complete MDM was able to support the transition of *Bd* from motile zoospores to cysts to germlings. In contrast to H-broth, *Bd* grown in MDM developed much more slowly and were unable to form zoosporangia (similar to **Figure 2C**).

### Effect of amino acids on Bd differentiation

The ability of cysts to progress into germlings was differentially affected by amino acid groups (**Figure 3A**). Removal of all amino acids from MDM resulted in a 10-fold decrease in encystment with concomitant loss of germling formation, which was indistinguishable from zoospores incubated in DSS. In contrast, the increase in encystment was not significantly decreased with the removal of any individual amino acid groups except for hydrophobic amino acids. Indeed, lack of hydrophobic amino acids resulted in a low abundance of germlings similar to the removal of all amino acids and the DSS control. Lack of basic amino acids also had a diminishing effect on germling formation, but it was not as pronounced as for hydrophobic amino acids.

**Figure 3.**
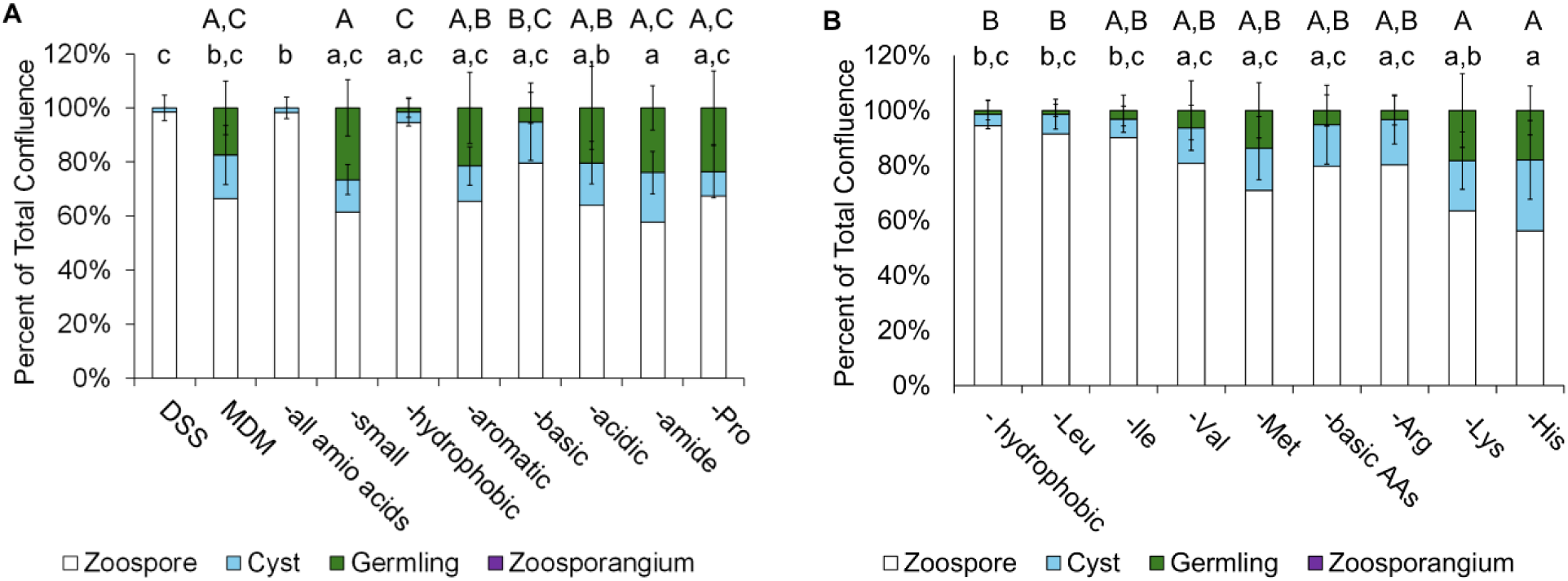
The effects of amino acids on *Bd* zoospore germination. **(A)** The effect of amino acids groups on *Bd* differentiation was determined by incubating zoospores (n=6 per condition) in MDM lacking either small (Ala, Ser, Thr, Gly, and Cys), hydrophobic (Leu, Ile, Met, and Val), aromatic (Phe, Tyr, and Trp), basic (Arg, Lys, and His), acidic (Asp and Glu), amide (Asn and Gln), or constrained (Pro) L-amino acids. **(B)** The effect of individual hydrophobic or basic amino acids on *Bd* differentiation was determined by incubating zoospores (n=6 per condition) in MDM lacking either Leu, Ile, Val, Met, Arg, Lys or His. Wells were imaged after 15 days of incubation at 19 °C, and the percentage of the confluence derived from zoospores (white sections), cysts (light blue sections), and germlings (green sections) were calculated. Groups labeled with the same letter are statistically the same (p>0.05). Lowercase letters indicate significance groups for cysts, uppercase letters indicate significance groups for germlings, and Greek letters indicate significance groups for zoosporangia.

To further test the effect of hydrophobic and basic amino acids on the encystment of zoospores and the germination of cysts into germlings, we individually removed L-Leu, L-Iso, L-Val, L-Met, L-Arg, or L-His from MDM (**Figure 3B**). As above, the level of encystment was reduced by the absence of L-Leu or L-Ile, and the germination into germlings was reduced by the absence of L-Leu, and to a lesser degree, by the absence of L-Ile, L-Val, or L-Arg (**Figure 3B**). Similarly, removal of basic amino acids showed that only L-Arg was necessary for robust germling formation.

### Effect of inorganic nitrogen on Bd differentiation

Since some chytrid fungi can utilize inorganic salts as sources of nitrogen^27^, we tested combinations of nitrate, nitrite, and ammonium salts paired with different counterions. As above, MDM (containing NH_4_HCO_3_ and Ca(NO_3_)_2_) only allowed encystment and germination into germlings (**Figure 4**). While the number of cysts present after 15 days of incubation varied among the different salt combinations, there appeared to be minimal effects on germling formation.

**Figure 4.**
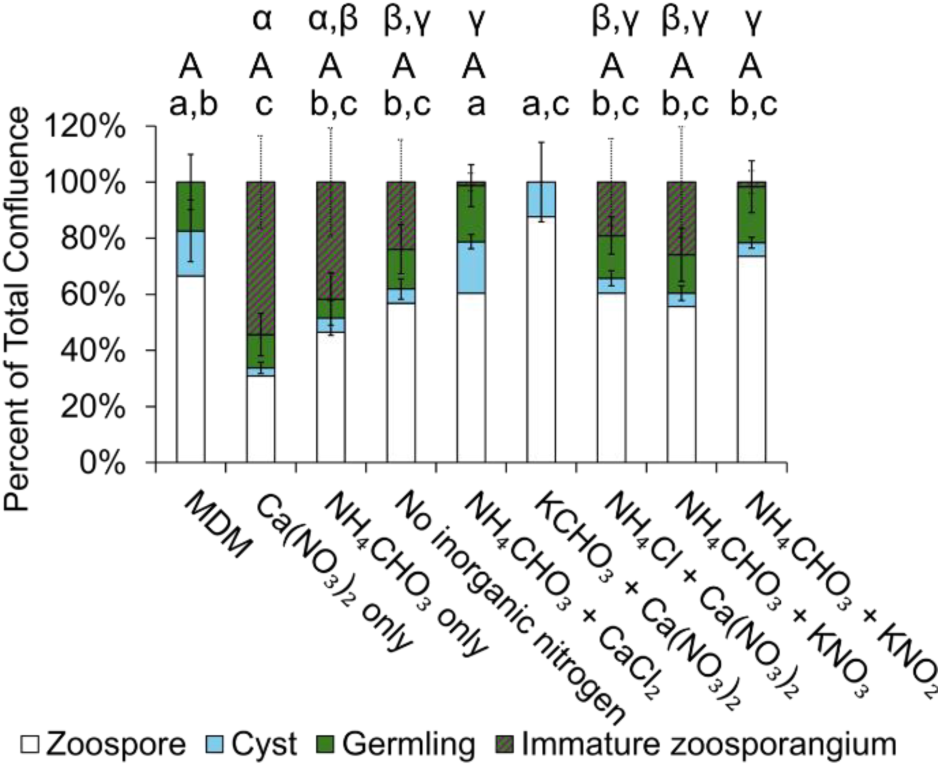
The effects of inorganic sources of nitrogen on *Bd* zoospore germination. The effect of inorganic nitrogen on Bd differentiation was determined by incubating zoospores (n=3 per condition) in MDM where Ca(NO_3_)_2_, and NH_4_CHO_3_ were substituted for single and binary combinations of Ca(NO_3_)_2_, NH_4_CHO_3_, KNO_3_, KNO_2_, CaCl_2_, KCHO_3_ and NH_4_Cl. Wells were imaged after 15 days of incubation at 19 °C, and the percentage of the confluence derived from zoospores (white sections), cysts (light blue sections), germlings (green sections), and immature zoosporangia (purple/green sections) were calculated. Groups labeled with the same letter are statistically the same (p>0.05). Lowercase letters indicate significance groups for cysts, uppercase letters indicate significance groups for germlings, and Greek letters indicate significance groups for zoosporangia.

Interestingly, whereas MDM only sustains the formation of germlings, removal of either NH_4_HCO_3_ or Ca(NO_3_)_2_ from MDM allowed further progression through the *Bd* lifecycle with significant production of immature zoosporangia. Removal of all inorganic nitrogen sources still resulted in zoosporangia formation, albeit at a lower level.

To test the inhibitory effect of the individual ions (NH_4_^+^, HCO_3_^-^, Ca^+2^, and NO_3_^-^) formed from the dissociation of these two salts, cations Ca^+2^ and NH_4_^+^ were substituted for K^+^, while anions HCO_3_^-^ and NO_3_^-^ were substituted for Cl^-^. Mixtures containing NH_4_HCO_3_/CaCl_2_ behaved similarly to complete MDM (NH_4_HCO_3_/Ca(NO_3_)_2_) in that no zoosporangia were formed. In contrast, mixtures containing NH_4_Cl/Ca(NO_3_)_2_ or NH_4_HCO_3_/KNO_3_ were able to form zoosporangia. To further test the requirements for zoosporangium formation, nitrate (NO_3_^-^) was substituted for nitrite (NO_2_^-^). This simple substitution abrogated the ability to form zoosporangia, with development stopping at the germling level, similar to zoospores grown in normal MDM.

### Effect of carbohydrates, vitamins, and trace minerals on Bd differentiation

In contrast to amino acids and inorganic nitrogen, neither carbohydrates, vitamins, or trace minerals affected the early stages of *Bd* differentiation. Interestingly, *Bd* zoospores were able to encyst and germinate into germlings in the absence of sugars (**Figure 5A**) at levels similar to MDM. Similar results were observed in the absence of vitamins (**Figure 5B**), or trace minerals (**Figure 5C**).

**Figure 5.**
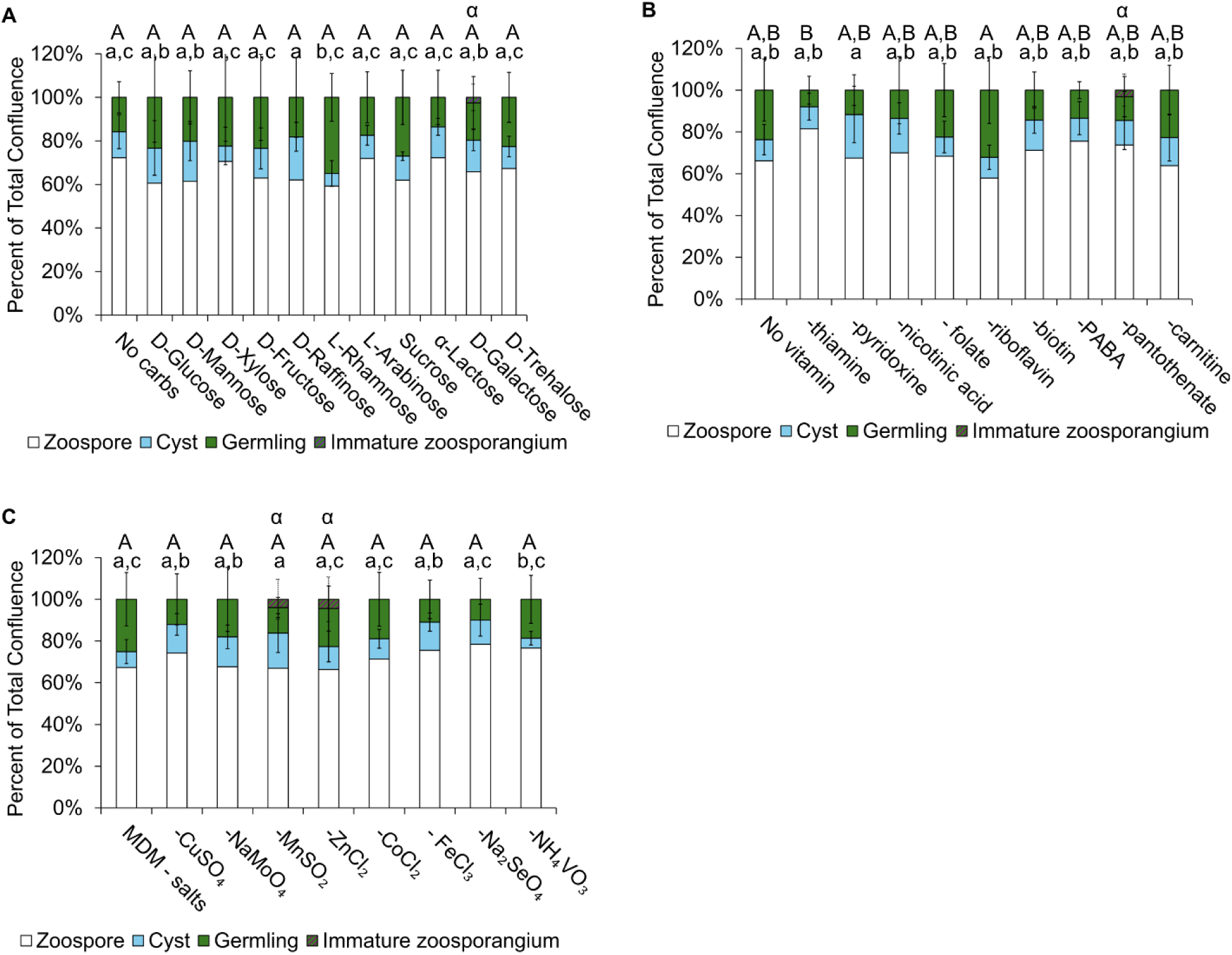
The effects of carbohydrates, vitamins, and trace minerals on *Bd* zoospore germination. **(A)** The effect of carbohydrates on *Bd* differentiation was determined by incubating zoospores (n=6 per condition) in MDM where D-glucose was substituted for other monosaccharides and disaccharides. **(B)** The effect of vitamins on *Bd* differentiation was determined by incubating zoospores (n=6 per condition) in MDM lacking specific vitamins. **(C)** The effect of minerals on *Bd* differentiation was determined by incubating zoospores (n=6 per condition) in MDM lacking specific trace minerals. Wells were imaged after 15 days of incubation at 19 °C, and the percentage of the confluence derived from zoospores (white sections), cysts (light blue sections), germlings (green sections), and zoosporangia (purple sections) were calculated. Groups labeled with the same letter are statistically the same (p>0.05). Lowercase letters indicate significance groups for cysts, uppercase letters indicate significance groups for germlings, and Greek letters indicate significance groups for zoosporangia.

### Effect of keratinized tissues on Bd differentiation

*Bd* can invade and degrade keratinized tissues^28^. Hence, we tested different keratinized tissues for their effect on the *Bd* lifecycle. Lizard scales, fish scales, and amphibian skin in DSS supported development into immature zoosporangia in the absence of any other nutrients (**Figure 6A**). Supplementation of DSS with bird feathers did not support progression of *Bd* zoospores past the encystment stage.

**Figure 6.**
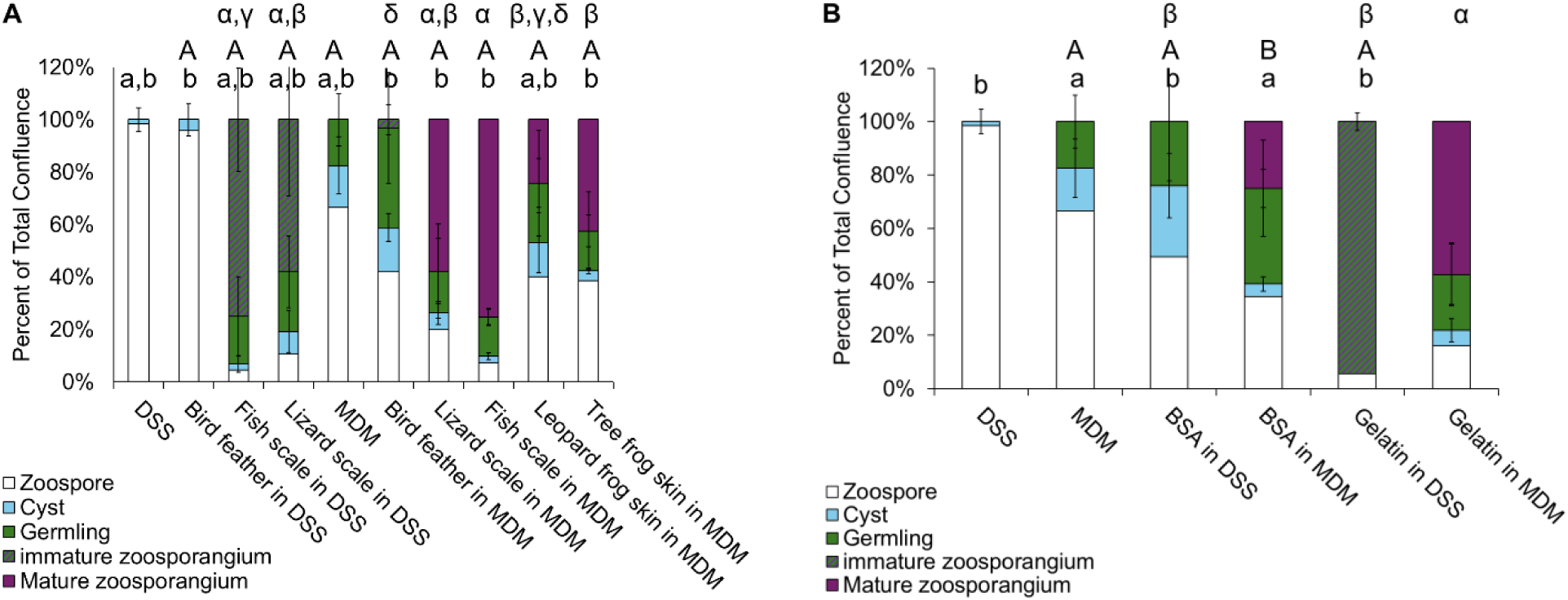
The effects of keratinized tissues and soluble proteins on *Bd* cell differentiation. **(A)** The effect of keratinaceous substrates on *Bd* differentiation was determined by incubating zoospores (n=6 per condition) in MDM or DSS supplemented with bearded dragon (*P. vitticeps)* scales, salmon (*O. nerka)* scales, pigeon (*C. livia*) feathers, relict leopard frog (*L. onca*) skin, or red-eyed tree frog (*A. callidryas*) skin. **(B)** The effect of proteinaceous substrates on *Bd* differentiation was determined by incubating zoospores (n=6 per condition) in MDM or DSS supplemented with either BSA or the soluble fraction of gelatin. All wells were imaged after 15 days of incubation at 19 °C, and the percentage of the confluence derived from zoospores (white sections), cysts (light blue sections), germlings (green sections), immature zoosporangia (purple/green sections), and mature zoosporangia (purple sections) were calculated. Groups labeled with the same letter are statistically the same (p>0.05). Lowercase letters indicate significance groups for cysts, uppercase letters indicate significance groups for germlings, and Greek letters indicate significance groups for zoosporangia.

When MDM was supplemented with reptile-, fish-, or amphibian-derived keratin sources, *Bd* was able to complete its lifecycle with the release of new zoospores from the keratin-sustained zoosporangia. Bird feathers did not have any effect on *Bd* differentiation in MDM.

Keratin sources have differential effects on the temporal development of *Bd* (**Table S1**). Supplementing MDM with *P. vitticeps* scales allows *Bd* zoospores to develop into cysts, reaching a maximum after 2-3 days. *Bd* cysts continue to develop into germlings after 4-5 days and convert to 50% mature, zoospore-producing zoosporangia at 7-15 days. Supplementing MDM with *O. nerka* scales follows a similar timeline as MDM supplemented with *P. vitticeps* scales and reaches up to 90% mature zoosporangia. In contrast, *C. livia* feathers only allow *Bd* to develop into germlings and no zoosporangia. Both of the amphibian skin keratin sources follow a similar development pathway, with *A. callidryas* creating double the number of zoosporangia compared to *L. onca*.

### Effect of soluble proteins on Bd differentiation

*Bd* grown in DSS supplemented with BSA did not progress beyond germlings (**Figure 6B**). In contrast, *Bd* grown in DSS supplemented with gelatin produced almost pure immature zoosporangia. Similar to keratinized tissues, when MDM was supplemented with either BSA or gelatin, *Bd* was able not only to form zoosporangia, but to release new zoospores.

When MDM was supplemented with BSA or soluble gelatin, zoospores developed with similar timeline through the formation of zoosporangia (**Table S2**). In contrast, supplementation of DSS with BSA results in the arrest of *Bd* development to the germling stage. Interestingly, supplementation of DSS with gelatin results in the accumulation of almost pure immature zoosporangia.

## Discussion

The ability of *Bd* to progress through its various life stages is important to establish infection. Notwithstanding the importance of *Bd* cell differentiation, previous attempts at identifying a suitable defined medium that can support *Bd* throughout its entire lifecycle have been unsuccessful^29^.

Our first challenge to study *Bd* cell differentiation was to develop a standardized method for quantifying the distinct *Bd* development stages. Because the diameters of different stages overlapped, using size as the single indicator for *Bd* cell cycle progression is insufficient. To adjust for size overlap, images of *Bd* cells were visually binned by both cell areas and the presence of cell-stage-specific structures. Specifically, if no cell showed the development of rhizoids, we concluded that *Bd* development stopped before getting to the germling stage. In this case, cells were categorized into a zoospore bin (diameter less than 5.0 μm) or a cyst bin (diameter greater than 5.0 μm). On the other hand, if any rhizoids were detected, cells were categorized into a zoospore bin (diameter less than 5.0 μm), a cyst bin (diameter between 5.0 μm and 6.0 μm), or a germling bin (diameter greater than 6.0 μm). Finally, if any enlarged germlings showed evidence of early zoospore formation (*e.g*., circular internal structures), then cells were categorized into a zoospore bin (diameter less than 5.0 μm), a cyst bin (diameter between 5.0 μm and 6.0 μm), a germling bin (diameter between 6.0 μm and 8.8 μm), or a zoosporangium bin (diameter greater than 8.8 μm).

To validate this binning system, we tested *Bd* development in the absence of nutrients (DSS) and in nutrient-rich media (H-broth) (**Figure 2**). DSS was previously developed as a medium that mimics the osmotic balance of pond or stream waters in which *Bd* is naturally found^31^. On the other hand, H-broth was used as a positive control for growth, as it has been established to support the complete lifecycle of *Bd*.

As previously reported^32^, a small population of zoospores can encyst without any nutrient but cannot progress any further, while remaining viable for up to two weeks (**Figure S1**). In agreement with previous studies^33,34^, we found that *Bd* grown in H-broth was able to complete their life cycle in 4-5 days (**Figure 2)**. Using our binning system, we were able to microscopically quantify *Bd* transformation from zoospores, to cysts, to germlings, to zoosporangia (**Figure 1**).

To test the effect of individual nutrients on *Bd* differentiation, we developed a minimal defined growth medium (MDM) that contained known quantities of amino acids, inorganic nitrogen sources, sugars, vitamins, and minerals. Since H-broth only contains tryptone and glucose, we expected that MDM would be sufficient to sustain the complete *Bd* life cycle.

Instead, zoospores could attach, encyst, and develop into germlings in the presence of MDM (**Figure 3A**) but could not progress through the maturation of zoosporangia and the release of new zoospores.

Since *Bd* encystment does not require nutrients, it was clear that one or more metabolites in MDM were required to complete the germination of cysts to germlings. Since tryptone present in H-broth is primarily composed of the digested peptides of casein, we first tested whether germling formation was affected by amino acids (**Figure 3)**. Indeed, we found that amino acids L-leucine and to a lesser extent L-isoleucine, L-valine, and L-arginine are required for the transition from cyst into germling.

It has previously been established that some chytrid fungi can utilize inorganic sources of nitrogen, often even as their sole source of nitrogen^27^. We observed a complex effect of inorganic nitrogens on *Bd* differentiation (**Figure 4**). Nitrate (NO_3_^-^), but not nitrite (NO_2_^-^), seems to be required for zoosporangium formation, even in the absence of proteinaceous sources. On the other hand, ammonium (NH_4_^+^) seems to be essential for germination of cysts into germlings and can also substitute for NO_3_^-^ in the formation of zoosporangia. The ability of NO_3_^-^ and NH_4_^+^ions to direct the formation of zoosporangia is strongly blocked by the combined action of ionic calcium (Ca^2+^) and bicarbonate (HCO_3_^-^). The inhibitory effects of Ca^2+^ and HCO_3_^-^ explain the inability of MDM to sustain *Bd* development pass the germling stage.

Because glucose and lactose are also major components in media used to grow *Bd*, we looked at the effects of carbohydrates in the *Bd* lifecycle (**Figure 5A**). Interestingly, we found germlings formation did not need carbohydrates. Similarly, we found no vitamin or trace mineral that was required for cell differentiation in *Bd* (**Figure 5B-C**).

The animal source of keratinized tissues had dramatic effects on *Bd* differentiation (**Figure 6A**). Amphibian skin, fish scales, and lizard scales in DSS were able to sustain the formation of zoosporangia. When supplemented with the metabolites of MDM, these same keratinized tissues allowed the completion of the *Bd* lifecycle by creating mature zoosporangia that released new spores. In contrast, feathers did not aid nor disrupted *Bd* differentiation.

To determine if the keratinized tissue enhancement of *Bd* differentiation was due to tissue degradation or simply attachment, we tested different sources of soluble proteins (**Figure 6B**). For this, we supplemented MDM and DSS with either BSA or with the soluble fraction of gelatin. Supplementation of DSS with BSA resulted in the formation of germlings in a manner similar to *Bd* grown in the presence of free amino acids, suggesting that BSA was getting degraded sufficiently to sustain the first stages of development. Soluble gelatin caused almost all *Bd* cells to progress to the immature zoosporangia stage, but not further. In the presence of other metabolites, both soluble gelatin and BSA behaved similarly to keratinized tissues in allowing completion of the *Bd* lifecycle and release of new zoospores.

Among keratin sources, lizard scales and fish scales enabled the greatest degree of maturation of zoosporangia (**Figure 6A**). However, reptile scales allowed for the completion of the first stages of *Bd* development more quickly but slowed down significantly during the transformation from germlings to zoosporangia (**Table S1**). In contrast, *Bd* cells grown in the presence of fish scales were able to produce zoosporangia more quickly. *Bd* cells grown in the presence of feathers, however, were unable to pass from the earliest developmental stages.

## Conclusions

In this study, we examined the effects of a variety of nutritional conditions on the ability of *Bd* to differentiate and complete its life cycle. As previously determined, zoospores did not need any nutritional signals to begin encysting, indicating that the transition from zoospore to cyst is not nutrient-gated.

We found that L-leucine, L-isoleucine and L-arginine were necessary for cysts to germinate into germlings. *Bd* zoospores have previously been shown to exhibit positive chemotaxis to glycine and L-cysteine^43^. In contrast, neither amino acid was required for *Bd* to encyst or germinate. This indicates that glycine and L-cysteine are not necessary for *Bd* to transition through its life cycle but are still specifically detected by zoospores.

The conversion of germlings to zoosporangia can be stimulated by two different mechanisms. On the one hand, this step could be activated by either ammonium or nitrate ions in the presence of amino acids and inhibited by calcium and bicarbonate. Alternatively, keratinized tissues and soluble proteins were sufficient for the formation zoosporangia in the absence of other nutrients. Neither of these pathways allowed for the maturation of zoosporangia. Finally, the release of new zoospores required the combined action of a protein/keratin source and a mixture of small molecule metabolites.

Carbohydrates serve a variety of unique and common roles in fungi. For example, mannose is a common cell wall carbohydrate^35^. Rhamnose is also a minor cell-wall component and a known requirement for pathogenicity in at least one plant-infectious fungus^36^.

Additionally, *Bd* has been shown to incorporate several monosaccharides, including glucose, mannose, galactose, and xylose into the extracellular matrix of cells involved in a biofilm^34^.

However, our results indicate that while carbohydrates still play important roles in *Bd* biology and metabolism, they are not essential signals for differentiation.

Some chytrids are known to grow normally in the absence of exogenous vitamins and may even be inhibited by high vitamin concentrations^37^. However, thiamine is often a required component of defined growth media for a variety of chytrid fungi^10,22^. Despite this, we found no evidence that the presence or absence of any of our tested vitamins, including thiamine, significantly affected the ability of *Bd* to develop to the germling stage.

*Bd* is known to infect only the keratinized epithelial tissue of its hosts by attaching to and degrading these structures into peptides and amino acids^38^. Furthermore, some microorganisms often utilize peptides more efficiently than amino acids^39^. In fact, *Bd* produces lytic enzymes that degrade various proteinaceous sources^28^. Indeed, we found that several keratin sources were able to signal *Bd* to progress to the later stages of its lifecycle, but the effects was dependent of the source of the keratinized tissue. These findings are of particular interest because reptiles^40^, fish^41^, and birds^42^ have all been proposed as potential vectors by which *Bd* has been able to spread between environments.

Finally, both keratin and gelatin more efficiently allowed for the development of *Bd* throughout its life stages compared to BSA. This is possibly because keratin and gelatin are natural targets for *Bd* when invading its host, whereas BSA is less biologically relevant to *Bd* and therefore less recognized as a substrate or nutrient.

Even though the rate at which *Bd* completed its life cycle was dependent on the protein/keratin source, it is unclear whether the protein sources tested were being metabolized or detected as signaling molecules. Further studies are being undertaken to elucidate these complex relationships.

As a corollary, we have identified nutritional conditions that allow for the arrest of *Bd* at various stages of its life cycle. Efficient cyst formation can be achieved by incubating zoospores in MDM containing KHCO_3_ and Ca(NO_3_)_2_. Although it was not possible to obtain germling cultures that did not contain cysts, MDM without riboflavin or with sucrose provided the largest proportion of germlings. In contrast, incubation of *Bd* with only gelatin produced an almost pure culture of immature zoosporangia. Finally, supplementation of MDM with a proteinaceous source allowed for zoospore release (**Figure 7**).

**Figure 7.**
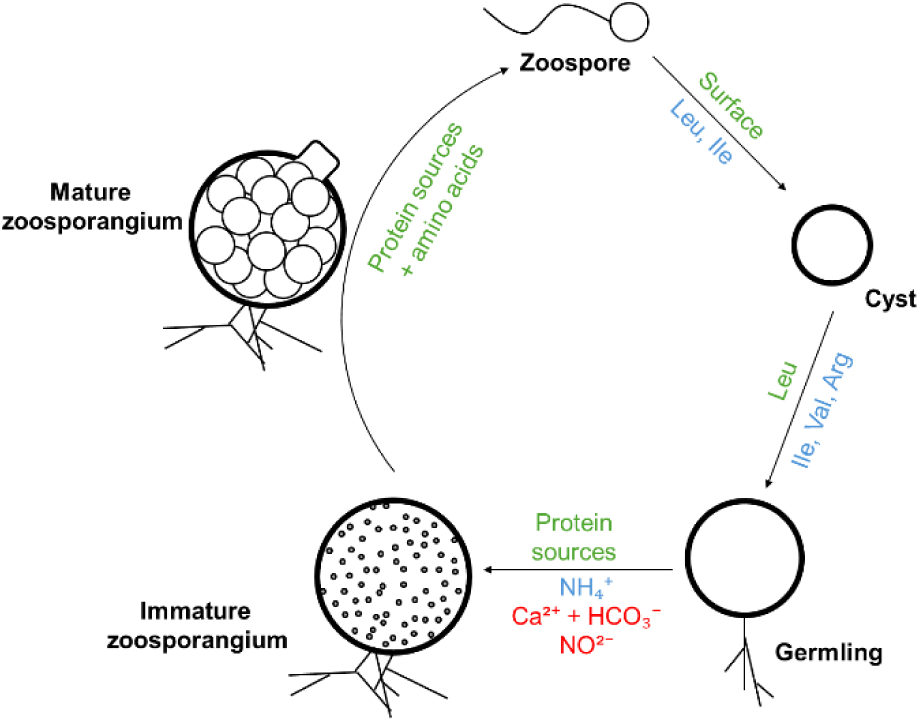
Nutritional scheme for *Bd* cell differentiation. Different cell stages are described in bold letters. Conditions in green are required for progression through developmental stages. Conditions in blue enhance progression through developmental stages. Conditions in red inhibit progression through developmental stages. *Bd* zoospores only require dilute salts for encystment. Similarly, L-leucine, L-isoleucine, L-arginine, and ammonium ions are required for cysts to develop into germlings. The formation of zoosporangia from cyst is stimulated by NO_3_^-^ and NH_4_^+^ ions, but their action is blocked by the combination of calcium and bicarbonate or nitrite alone. Cyst differentiation into zoosporangia can also be induced by some keratin and soluble protein sources in the absence of other nutrients. Finally, the maturation of zoosporangia and release of new zoospores is stimulated by the combined action of protein sources and amino acids.

## Acknowledgements

This work was supported by the National Institute of Health [grant numbers R01-AI109139 and]. We want to thank Dr. Jef Jaeger for donation of *Bd* strain LBP114, and Josh Levy and Rebeca Rivera for assistance in culturing methods. We would also like to acknowledge Snow the bearded dragon Okada, Tikki the pigeon, Harold the red-eyed tree frog, and Ferdinand the bull frog for contributing keratinized tissues to this work. We also thank Dajahneek O’Brien, Dinh Huynh, Leanna Oliver, and Erica Bacab for assistance in image processing.

